# IL-8 Instructs Macrophage Identity in Lateral Ventricle Contacting Glioblastoma

**DOI:** 10.1101/2024.03.29.587030

**Authors:** Stephanie Medina, Asa A. Brockman, Claire E. Cross, Madeline J. Hayes, Bret C Mobley, Akshitkumar M. Mistry, Silky Chotai, Kyle D Weaver, Reid C Thompson, Lola B Chambless, Rebecca A. Ihrie, Jonathan M. Irish

## Abstract

Adult IDH-wildtype glioblastoma (GBM) is a highly aggressive brain tumor with no established immunotherapy or targeted therapy. Recently, CD32^+^ HLA-DR^hi^ macrophages were shown to have displaced resident microglia in GBM tumors that contact the lateral ventricle stem cell niche. Since these lateral ventricle contacting GBM tumors have especially poor outcomes, identifying the origin and role of these CD32^+^ macrophages is likely critical to developing successful GBM immunotherapies. Here, we identify these CD32^+^ cells as M_IL-8 macrophages and establish that IL-8 is sufficient and necessary for tumor cells to instruct healthy macrophages into CD32^+^ M_IL-8 M2 macrophages. In *ex vivo* experiments with conditioned medium from primary human tumor cells, inhibitory antibodies to IL-8 blocked the generation of CD32^+^ M_IL-8 cells. Finally, using a set of 73 GBM tumors, IL-8 protein is shown to be present in GBM tumor cells *in vivo* and especially common in tumors contacting the lateral ventricle. These results provide a mechanistic origin for CD32^+^ macrophages that predominate in the microenvironment of the most aggressive GBM tumors. IL-8 and CD32^+^ macrophages should now be explored as targets in combination with GBM immunotherapies, especially for patients whose tumors present with radiographic contact with the ventricular-subventricular zone stem cell niche.

- IL-8 is expressed by GBM cells and enriched in lateral ventricle-contacting tumors
- M_IL-8 macrophages are CD32^+^ HLA-DR^++^ CD163^+^ CD206^+^ CD86^-^ PD-L1^-^
- M_IL-8 macrophages instructed with IL-8 or GBM conditioned medium match human glioblastoma associated macrophages
- IL-8 is necessary for GBM tumor cells to generate M_IL-8 macrophages

## Introduction

Glioblastoma (GBM) is the most aggressive and most common primary brain tumor among adults, with a poor prognosis of less than 2 years ^1–3^. While immunotherapies have revolutionized the treatment of many solid tumors including melanoma, lung, and kidney cancer, GBM tumors remain resistant to immunotherapies, in part due to an immune tumor microenvironment (TME) that is refractory to these agents ^4,5^.

Glioblastoma-associated macrophages and microglia are the most abundant immune cell type infiltrating tumors and can often represent up to half of all the cells of the tumor mass ^6–10^. The majority of GBM tumor macrophages are presumed to originate from circulating blood monocytes ^11^. Upon entry into the TME, these tumor macrophages adopt new cellular and molecular identities that are critical for GBM progression. Macrophages can contribute to immune suppression through the expression of surface proteins like CD163, a marker of immunosuppressive macrophages that contributes to T cell dysfunction and has been associated with poor prognosis in GBM ^12–14^. High dimensional immune cell profiling in GBM recently identified a subset of CD32^+^ HLA-DR^hi^ monocyte derived macrophages that were distinguished by a potentiated response to inflammatory cytokine signaling via p-STAT3. The abundance of CD32^+^ GBM associated macrophages (GAMs) independently stratified patient survival, and these GAMs predominated in the microenvironment of tumors that contacted the lateral ventricles, the location of the ventricular-subventricular zone (V-SVZ) stem cell niche ^15^. V-SVZ contact by GBM tumors is established as closely associated with poorer clinical outcomes for patients^16,17^. Reprograming immunosuppressive GAMs into more inflammatory states has been shown to have potential clinical benefit ^18–20^. Thus, understanding the mechanisms that generate CD32^+^ GAMs in V-SVZ-contacting human tumors could provide a strategy to rehabilitate the immune microenvironment of the most aggressive subtype of a deadly brain tumor.

Tumor associated macrophages play diverse roles in cancer development and tumor progression leading to poor prognosis of many solid tumors beyond GBM ^21^, such as breast ^22^, head and neck ^23^, bladder^24^, melanoma^25^, and prostate cancer ^26^. Thus far, murine models and patient derived xenograft models have been used to understand the heterogeneous phenotypic states of infiltrating microglia and macrophages in GBM ^27–31^. However, these models may not reflect all aspects of human immunology and may not reflect structural features of the human brain or GBM tumors. For example, *CXCL8*, the gene for human IL-8 protein, is one of the 20% of human genes that lacks a mouse ortholog. It is vital to continuously improve preclinical models, and one area for urgent attention is to ensure that the immune microenvironment of different models closely reflects that observed in human tumors.

Historically, healthy macrophage identity has been presented as aligned to one of two extremes: M1-like macrophages that promote an inflammatory immune response or M2-like macrophages that promote a suppressive immune response ^32^. However, the last decade of research has revealed and characterized a spectrum of macrophage identities that can be tracked through surface proteins which are closely linked to diverse functions and activation states ^33,34^. Current state of the art approaches define macrophage activation states based on surface immunophenotype and name macrophage states based on the cytokines leading to their specialization. For example, M1-like, IFNγ polarized macrophages (M_IFNγ) are distinguished by elevated expression of CD86, a surface protein that provides costimulatory signals promoting T cell activation and survival ^35^, PD-L1 ^36^, the programed cell death ligand receptor, and lack of expression of scavenger receptor CD163 and mannose receptor CD206 ^33,37^. In contrast, M2-like macrophages can include those polarized by IL-6 or by IL-4 (M_IL-6 or M_IL-4). These macrophage activation states both express high levels of surface CD163 and CD206 proteins, but M_IL-6 express higher levels of the FCγRII CD32 ^34,38^. In addition, myeloid derived suppressor cells (MDSCs) are distinguished functionally by their ability to suppress T cell proliferation and are characterized by low expression of cell surface HLA-DR protein^39^.

The tumor microenvironment can produce cytokines/chemokines, which are involved in the recruitment of normal cells to promote growth, invasion, angiogenesis, and metastasis of glioblastoma. The exact cytokines that are secreted from glioblastoma tissue may vary depending on the specific case and the stage of the disease^40^. While it is established that macrophages can respond distinctly to select stimuli in their environment^32^, other cytokines, such as IL-8, have not been studied as extensively in the context of macrophage polarization. Therefore, it is not clear whether a monocyte derived M_IL-8 cell exists, whether it ‘leans’ to M1 or M2, and whether it is phenotypically or functionally distinct from M_IL-6 macrophages or other subtypes.

IL-8 was originally described as a chemokine whose main functions are generally reported to be attraction of neutrophils via receptors CXCR1 and CXCR2 ^41,42^. It is now known to play key roles in wound healing, angiogenesis, inflammation, and tumor growth ^43,44^, and it has been most studied in cancer in the context of epithelial origin tumors^45^. In GBM, IL-8 was reported to be highly expressed and to play a role in regulating GBM stem cells, promoting tumor growth, and promoting angiogenesis ^46–49^. Previous work focused on IL-8’s ability to activate vascular mimicry in tumor cells, including after treatment with alkylating chemotherapy temozolomide ^50^. Il-8 has been negatively correlated with glioma patient survival ^51,52^, but has not previously been linked to macrophages in GBM or to the aggressive subtype of GBM that presents in contact with the V-SVZ.

Here, we use *ex vivo* culture of primary GBM tumor cells and healthy blood derived macrophages to model the generation of suppressive CD32^+^ GAMs. IL-8 is identified as the primary factor directing macrophage identity in human GBM tumors, establishing a novel role for IL-8 in the tumor microenvironment and in GBM.

## Results

### Healthy blood macrophages polarized *ex vivo* by IL-6 contrast with macrophages in human GBM

In this study, GBM tumor conditioned medium (**<u>GBM_TCM</u>**) was added to the field-standard macrophage polarization assay used to generate inflammatory and suppressive macrophages from healthy monocytes ^37,53^. The goal of this experiment was to establish whether macrophages polarized with GBM_TCM (**<u>M_GBM_TCM</u>**) closely resemble those observed *in vivo* in human GBM tumors (GBM-associated macrophages, **<u>GAMs</u>**). GBM_TCM for macrophage polarization was created by culturing dissociated single cells from primary human glioblastoma tumors for 3 days (see Methods and ^37^ for additional detail). After 3 days of culturing monocytes in the presence of macrophage colony stimulating factor (M-CSF), the resulting macrophages were stimulated with GBM TCM for an additional 3 days, analyzed using spectral flow cytometry, and their phenotype compared to GAMs or to canonical healthy macrophage subtypes representing M1 (M_IFNγ) and M2 (M_IL-6).

**Box 1.**
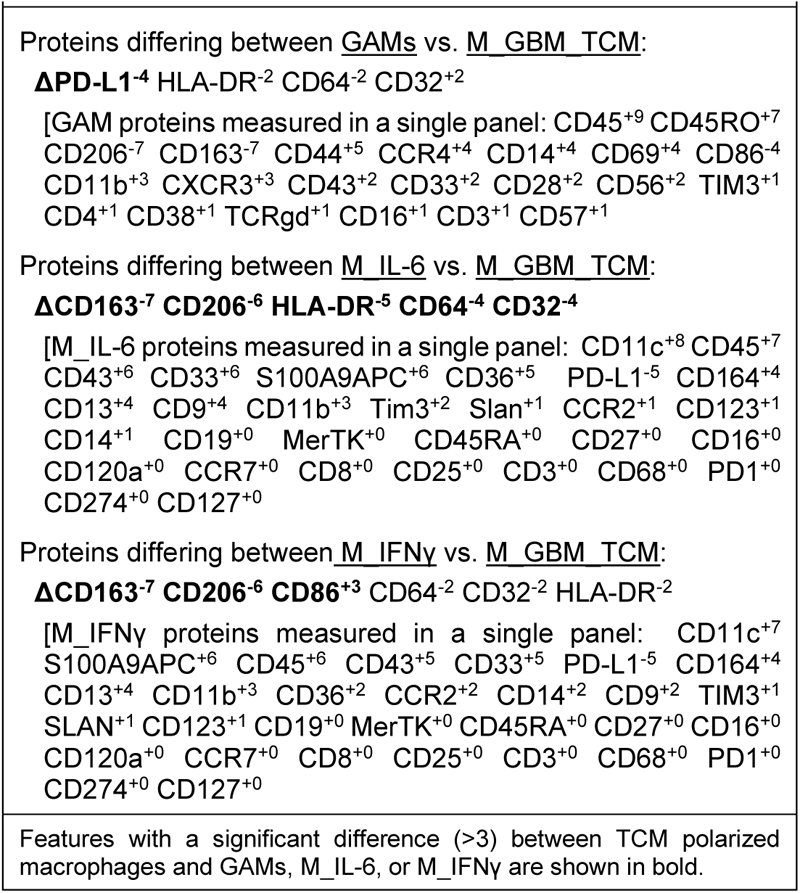
Quantitative comparison of proteins expressed on GAMs, M_IL-6, or M_IFNγ to M_GBM_TCM macrophages using MEM.

To determine whether M_GBM_TCM macrophages model the generation of previously described GAMs, we performed Marker Enrichment Modeling (MEM) analysis ^37,54^ using published cytometry data from primary human GBM macrophages ^15^ and newly generated data from macrophages subjected to different stimulatory conditions, including TCM, M2 cytokines IL-6 and IL-4, and M1 control cytokine IFNγ ^33^. Following MEM, a ΔMEM analysis ^55^, which subtracts two MEM labels to identify differentially enriched features, was performed to quantify and compare changes in protein expression between macrophages. Published cytometry panels and newly generated data panels shared 4 features that we compared in this analysis: CD64, CD32, HLA-DR/MHC II, and PD-L1. Of these, CD32 and HLA-DR were signature features that distinguished C-GBM GAMs, and the classic M1 feature PD-L1 was observed to be missing on C-GBM GAMs. The newly generated data also measured CD86 (M1), CD206 (M2), and CD163 (M2) macrophage markers. The ΔMEM analysis showed that TCM produces cells that differ from IL-6 and other conditions but were a close match for GAMs observed in C-GBMs. While M_GBM_TCM were similar to GAMs in expression of HLA-DR, CD64, and CD32 proteins, GAMs generally had lower PD-L1 expression compared to M_GBM_TCM (**Box 1**). In contrast, IL-6 or IFNγ stimulations did not lead to as much CD163 or CD206 expression as TCM. IL-6 did not trigger expression of CD32, HLA-DR, or CD64 and IFNγ led to much higher expression of CD86 than TCM (**Box 1**). In addition to the markers shared between panels, MEM was used to evaluate features enriched or missing from these macrophage populations (Box 1, proteins measured in a single panel). Taken together, these results indicated that IL-6 and other tested cytokines alone were not sufficient to generate macrophages with the phenotype observed *in vivo* in GBM tumors.

### Healthy blood macrophages polarized *ex vivo* with tumor conditioned media are CD32^+^ M2 cells comparable to primary GBM macrophages

Having established key features expressed by macrophages generated with GBM TCM, the goal was next to understand percent positivity for different proteins using a traditional gating strategy. Using manual expert gating, polarized macrophages were analyzed to quantify the amount of CD206 and CD163 positive cells, CD32 and CD163 positive cells, CD163 positive and CD86lo expressing cells, and CD163 negative and PD-L1hi expressing cells (**Figure 1A**). Unpolarized monocytes expressed low levels of all markers and contained less than 5% of cells positive for any markers assessed. Less than 12% of M_IFNγ macrophages were found to be CD163^+^CD32^+^CD206^+^ and CD86^lo^. However, over 80% of M_ IFNγ macrophages were gated as CD163^-^ PD-L1^hi^ cells, which is expected for M1 macrophages. M_IL-6 and M_GBM_TCM macrophages consistently expressed M2 phenotype markers: over 39% of cells were CD206^+^ CD163^+^, over 34% of cells were CD163^+^CD32^+^, over 26% of cells were CD163^+^ CD86^lo^, and less than 1% were CD163^-^ PD-L1^hi^ for all conditions (**Figure 1B**). This established that in addition to phenocopying GAMs, M_GBM_TCM macrophages exhibit a suppressive M2 like phenotype.

**Figure 1.**
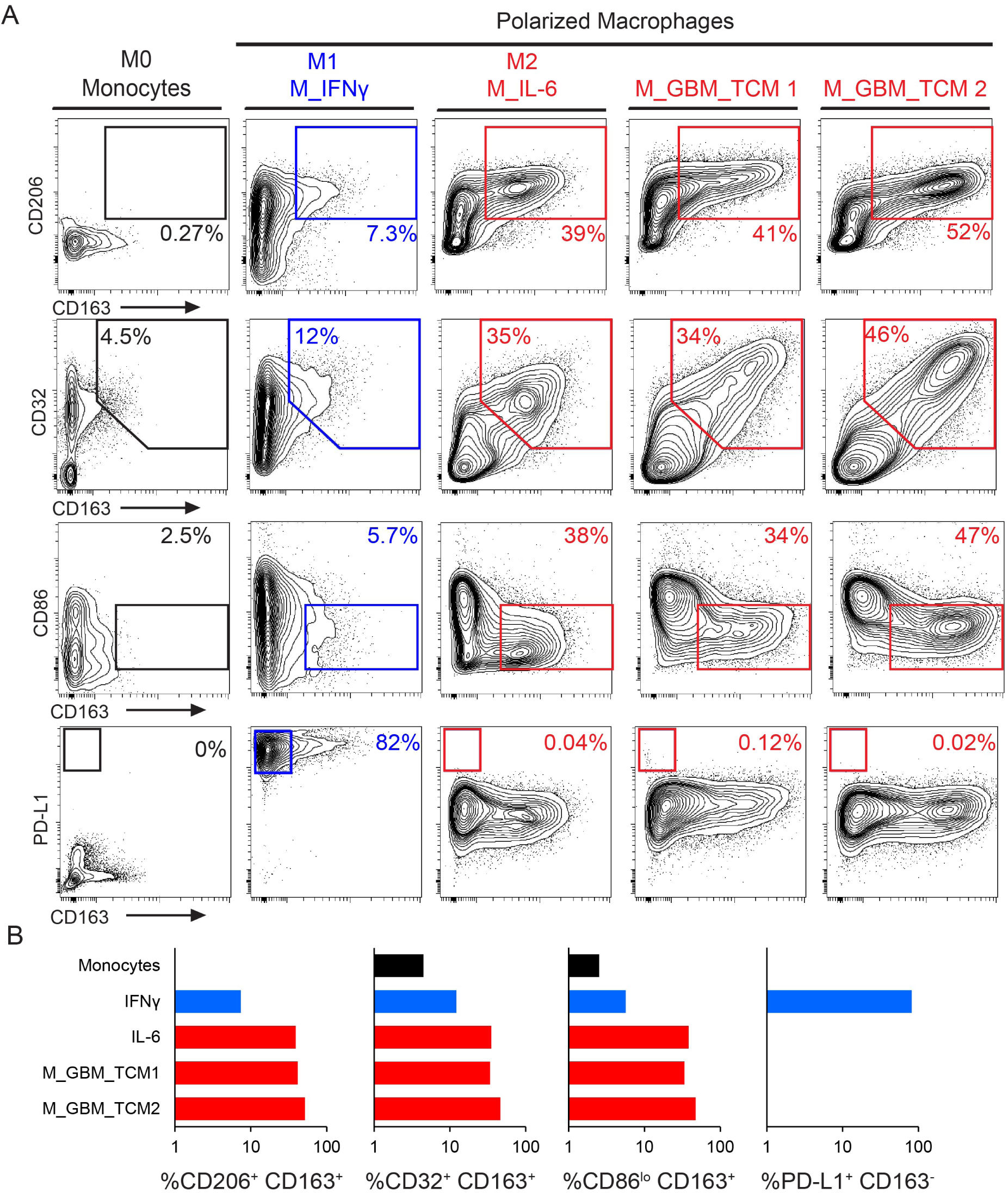
M_GBM_TCM are similar to an M2 phenotype and distinct from other M2 macrophages, including M_IL-6. A) 2D contour plots show expression of signature surface proteins CD163, CD206, CD86, and PD-L1 for each condition. Cells were either unstimulated monocytes, macrophages stimulated with IFNγ (a control for M1-like macrophages), macrophages stimulated with IL-6 (a control for M2-like macrophages), or macrophages stimulated with primary tumor conditioned media collected after 3 days of ex vivo culture (sample ID: TCM1, TCM2). B) Bar graphs display the percentage of cells included in indicated gates across conditions.

### IL-8 was produced by cells from all tested primary human GBM tumors

To learn more about potential mechanisms by which M_GBM_TCM macrophages are polarized into a suppressive phenotype it was important to dissect the factors that are secreted by tumor cells during *ex vivo* culture. The goal of this experiment was to identify any potential candidate mediators of macrophage polarization in GBM. To identify secreted proteins present in GBM TCM, an array analysis testing for 105 different soluble factors and cytokines was performed on TCM from 7 primary human tumor samples (**Figure 2**). Unconditioned media was used as a negative control (**Figure 2A**). Quantification of dot intensity density, which is relative to the quantity of protein present in any sample, revealed 6 potential candidates whose median secretion was above the calculated threshold of significance across all 7 tumors: Emmprin, CXCL8/IL-8, Macrophage migration inhibitory factor (MIF), Matrix metalloproteinase-9 (MMP9), Osteopontin, and Serpin E1 (**Figure 2B**) Only IL-8 was consistently secreted at a high level by cells from all tested GBM tumors (**Figure 2B**, dark blue dots, N=7).

**Figure 2.**
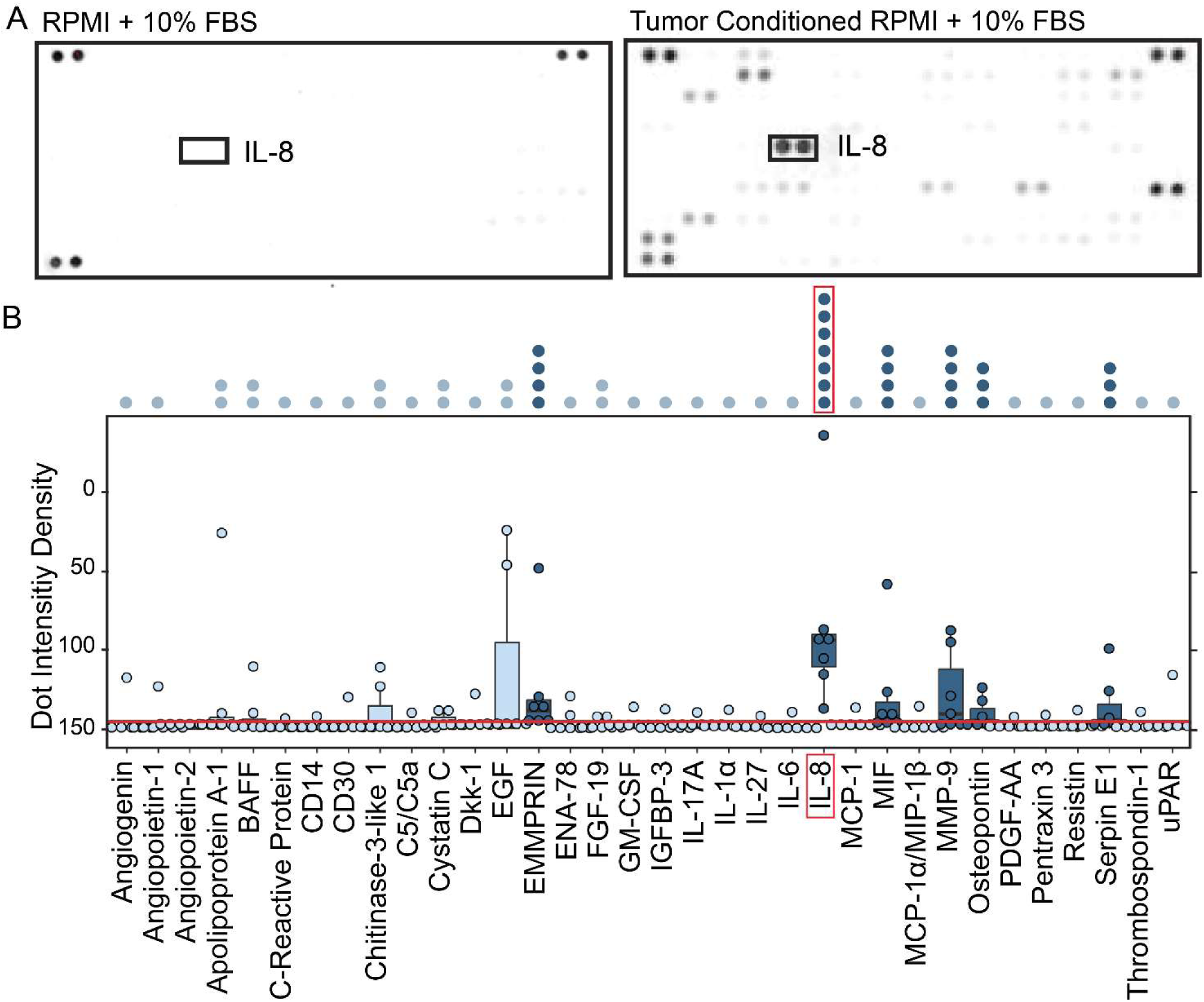
IL-8 is secreted by glioblastoma tumor cells *ex vivo*. A) Cytokine arrays are shown from media negative control (RPMI) and media conditioned by a representative glioblastoma tumor (RPMI + LC-26). Darkness of the spot indicates increasing presence of one of the 105 tested cytokines. Positive and negative control spots are located on each corner of the blot. B) Box plot shows quantification of the intensity density of proteins measured on the cytokine arrays of GBM tumor conditioned media (N = 7 tumors). A significance threshold was calculated based on three standard deviations above the level of background observed in negative control wells (red line). Cytokines present in at least 1 of 7 tumors are labeled across the X axis. Significant cytokines where the median expression across tumors exceeded the calculated threshold are colored in dark blue. Non-significant cytokines are colored light blue. Stacked dots above the plot indicate the number of tumor samples that exceeded the threshold for any cytokine measured. The most significant cytokine identified is labeled by a red box. The full dataset is available online (Supplementary Information).

### Blocking IL-8 abrogates M2 macrophage polarization by GBM tumor cells

After identifying IL-8 as the most abundant and reproducible cytokine secreted into TCM across tumors, we hypothesized that IL-8 is necessary and sufficient to generate CD32^+^ M_GBM_TCM macrophages *ex vivo.* To test this hypothesis, macrophages were polarized over 3 days in culture in the presence of either recombinant IL-8 or GBM TCM and with or without a monoclonal blocking antibody against IL-8 (α-IL-8), which remained present throughout macrophage polarization. Using spectral flow cytometry, the expression of signature M_GBM_TCM surface markers was measured.

To gain a visual understanding of how macrophage identity shifts across different polarization conditions, a t-SNE analysis of all polarized macrophages based on all measured surface marker features was performed. t-SNE plots displaying the cellular density across the map revealed that both M_IL-8 and M_GBM_TCM macrophage conditions lacking α-IL-8 predominantly clustered towards the top left of the t-SNE map (>57%). Macrophages polarized in the presence of an IL-8 blocking antibody (+ α-IL-8) were found to predominately cluster towards the bottom right of the t-SNE map (>58%) (**Figure 3A**).

**Figure 3.**
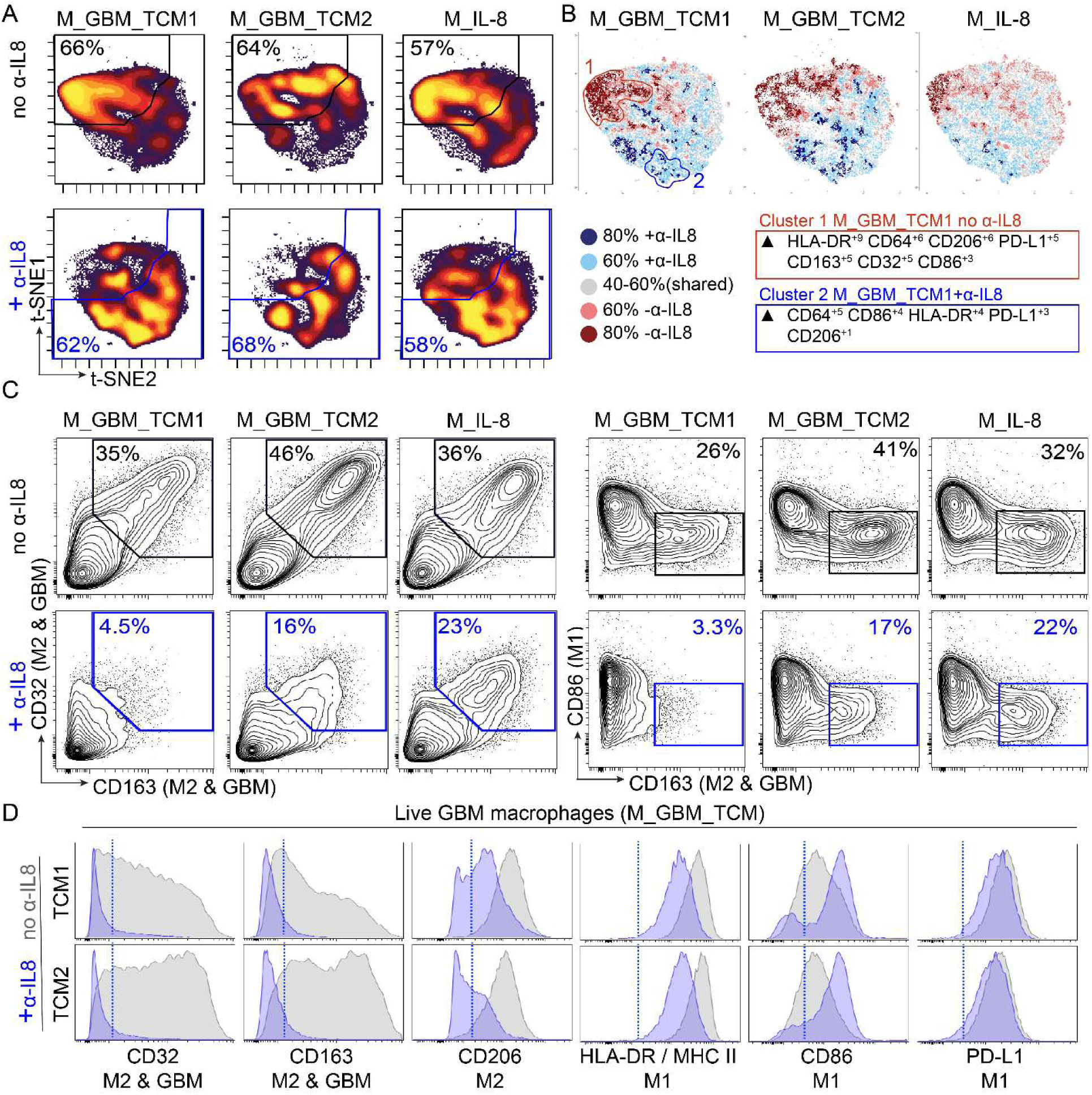
GBM secreted IL-8 mediates *ex vivo* polarization of macrophages to express a suppressive signature. A) t-SNE plots display cell density of polarized macrophages across different conditions including recombinant IL-8 (M_IL-8) and GBM tumor conditioned media (M_GBM_TCM), in the presence or absence of α-IL-8 blocking antibody. B) T-REX analysis comparing polarization conditions in the presence or absence of α-IL-8 where cells that are >80% enriched in macrophages polarized in the absence of α-IL-8 are colored in dark red, >60% enriched in light red, cells enriched >80% in macrophages polarized in the presence of α-IL-8 are colored in dark blue and >60% enriched in light blue. Cells colored in gray are similarly enriched in both conditions C) 2D contour plots show expression of signature surface proteins CD163, CD32, CD86, and PD-L1 for each condition. Gates colored in blue indicate the conditions where macrophages were polarized in the presence of α-IL-8. D) Histogram plots display individual protein expression of markers CD32, CD163, CD206, HLA-DR, CD86 and PD-L1. Overlaid histograms represent macrophages polarized in the presence (blue) or absence (gray) of α-IL-8 blocking antibody. A blue dotted line indicates a threshold for positive expression.

To quantify differences in phenotypes, a T-REX analysis ^56^ was performed to compare the macrophage phenotypes that were generated in the presence or the absence of α-IL-8 (**Figure 3B**). T-REX plots revealed condition-specific clusters of macrophages enriched when macrophages were polarized in either the presence (colored in blue) or absence (colored in red) of α-IL-8. MEM analysis of these condition specific clusters revealed that cells in, cluster 1 which were enriched in conditions lacking α-IL-8 expressed higher levels of HLA-DR, CD206, CD163, and CD32 but lower levels of CD86 in comparison to cells in cluster 2 which were enriched in conditions containing α-IL-8.

To get a closer look at specific expression of GAM signature proteins, expert gating was used to quantify CD163^+^ CD32^+^ CD86^lo^ cells in conditions with or without the addition of α-IL-8 (**Figure 3C**). M_IL-8 and M_GBM_TCM macrophages displayed over 35% positivity CD163^+^ and CD32^+^ cells, and over 26% positivity for CD163^+^CD86^lo^ cells. Upon addition of α-IL-8 blocking antibody, the percentage of CD163^+^ and CD32^+^ and CD163^+^CD86^lo^ was reduced to under 23% for M_IL-8 cells and under 17% of cells for M_GBM_TCM conditions.

Finally, histograms were used to view the effects of α-IL-8 on the expression of individual protein markers (**Figure 3D**). This highlighted how blocking IL-8 in TCM prevents the expression of M2/GAM markers (CD32 and CD163) in polarized macrophages. In addition, blocking IL-8 promotes a higher expression of M1 marker CD86, but has little to no effect on PD-L1 expression. These results support two major findings: 1) IL-8 on its own is sufficient to polarize macrophages towards a phenotype that is similar to M_GBM_TCM, and IL-8 in GBM TCM is necessary for the polarization of M_GBM_TCM macrophages.

### IL-8 is common in primary tumors and more highly expressed in C-GBM tumors

To confirm that IL-8 secretion also occurs *in vivo* in primary human GBM tumors, immunohistochemical (IHC) staining for CXCL8 (IL-8) was performed on a tumor microarray (TMA) composed of primary tumor samples from 73 IDH-wt GBM patients. To evaluate IL-8 expression across primary GBM tumor samples, the percentage of IL-8 positive pixels per cell and per core was quantified for every patient core (3 cores/patient, 73 patients total). A wide range varying from 0%-50% positive pixels of IL-8 signal were detected across cells and patients (**Figure 4 A-B**). Next, to evaluate IL-8 expression patterns across tumor tissue, cellularity per core was quantified and plotted against % positive pixels. In addition, visual validation confirmed IL-8 staining patterns were predominantly extracellular as opposed to nuclear or cellular (**Figure 4C**). This confirmed that IL-8 secretion is a feature that was conserved from *in vivo* tumors in the *ex vivo* GBM model system.

**Figure 4.**
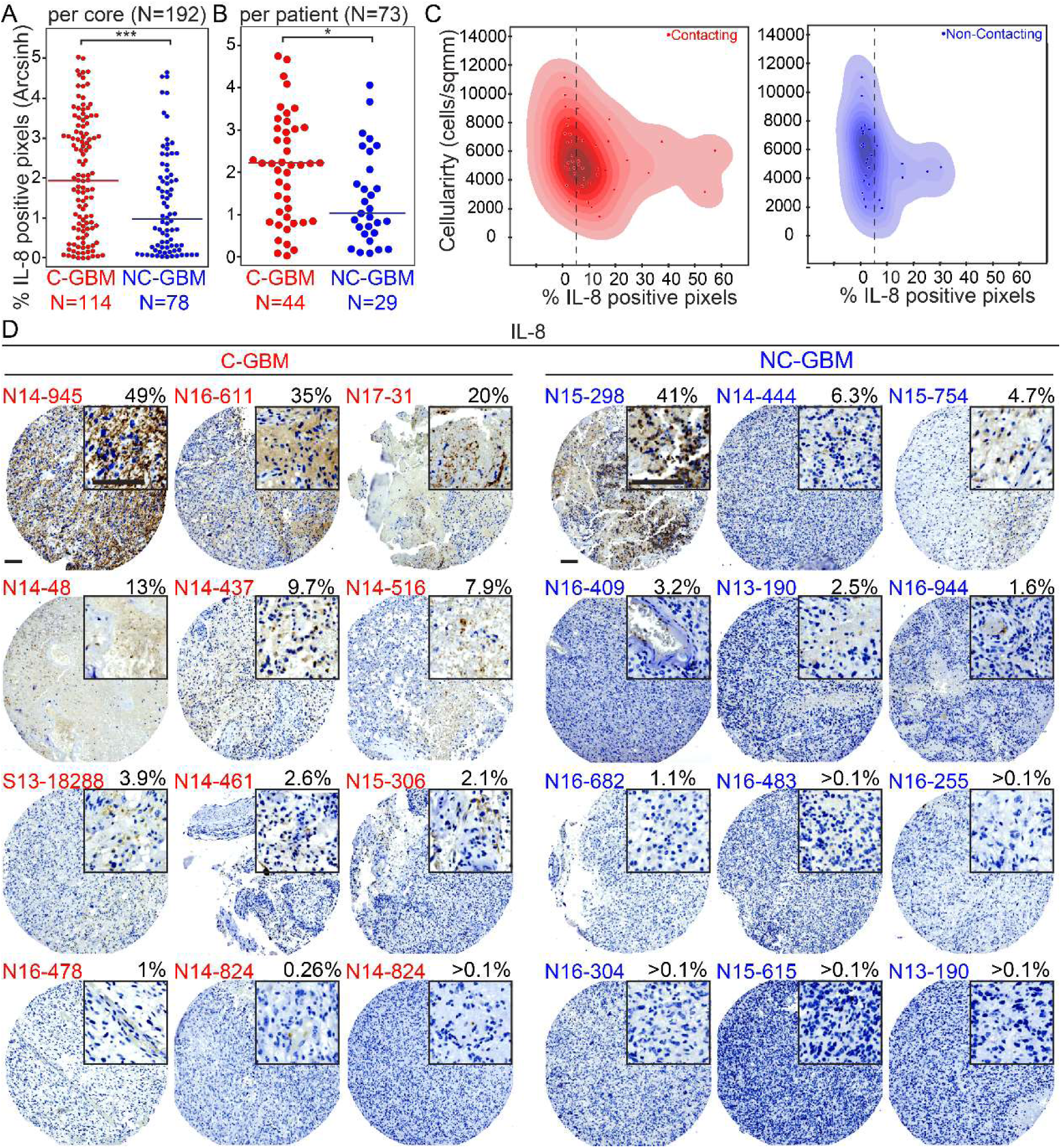
IL-8 is expressed in primary contacting tumors and less abundantly expressed in non-contacting tumors. A) Dot plot comparing the percentage of positive pixels per core stained for IL-8 in contacting (N=114) and non-contacting (N=78) GBM tumor cores. 3 cores of each patient sample were included for a total of 192 cores. A Mann Whitney statistical test was used to analyze the difference in immunohistochemical staining of IL-8 between the two groups. *** indicates p < 0.001. B) Dot plot graph comparing the average percentage of positive pixels per patient (N=73) stained for IL-8 in contacting (N=44) and non-contacting (N=29) GBM tumors. A Mann Whitney statistical test was used to analyze the statistical difference in immunohistochemical staining of IL-8 between the two groups. * Indicates p < 0.05. Dot plot comparing number of cells per mm^2^ (on an arcsinh scale) to percentage of positive IL-8 pixels for each patient. Contacting tumors are displayed in red and non-contacting tumors are displayed in blue. Representative immunohistochemistry images of CXCL8 (IL-8) expression on glioblastoma tumor microarray (TMA) (12 Contacting tumor examples shown with red labels and 12 Non-Contacting tumor examples shown with blue labels). Top right corner of each image contains 3x zoomed in view (scale bar=100µm).

Presuming that M_GBM_TCM macrophages model CD32+ GAMs previously described as a distinguishing immune cell subset for tumors that contact the V-SVZ (C-GBM), we hypothesized that if IL-8 is necessary for the generation of M_GBM_TCM macrophages, then IL-8 would be more abundantly expressed in C-GBM. To test this hypothesis, every tumor sample included in the TMA was radiographically scored and classified as contacting (C-GBM) or non-contacting (NC-GBM). The percentage of positive pixels per core was quantified and compared between C-GBM cores and NC-GBM cores, revealing that IL-8 was more significantly expressed in C-GBM tumors (p<0.001, N=73) (**Figure 4C**).

Images of representative cores were then chosen to determine whether IL-8 protein expression was primarily overlapping with non-immune tumor cells or immune cells. Strikingly, when IL-8 was expressed at high levels, it was apparently present in non-immune tumor cells (**Figure 4**). Taken together, these results indicate that in human GBM tumors that contact the V-SVZ, GBM cells produce IL-8 which shifts incoming macrophages into CD32^+^ M2 macrophages highly similar to M_IL-8s produced *ex vivo* by IL-8 and TCM.

## Discussion

Immune cells are known to play a critical role in the development and the progression of tumors. Thus, tumor immune evasion has become recognized as a hallmark of cancer ^57,58^. The modulation of immune cells represents one of the main driving features of GBM^59^, and GBM subtypes have been shown to modulate their TME through the aberrant secretion of multiple factors^60^, especially including those that contribute to immunosuppression^61^. Myeloid derived macrophages, brain resident microglia, dendritic cells, and myeloid derived suppressor cells (MDSCs) are the main components of the GBM TME^60^. Since macrophages are known to respond to stimuli in their microenvironment, we hypothesized that a GBM secreted soluble factor might play a role in instructing the aggressive macrophages that have been shown to drive the immunosuppressive microenvironment of GBM. We predict that these suppressive macrophages may be functioning at the end of the cancer immunity cycle by suppressing T cell activities that would ultimately lead to cancer cell death^58^. The focus of this study was to understand how GBM tumor secreted factors instruct the functional identity of previously described, survival stratifying, immunosuppressive GAMs. In this study, an *ex vivo* cell culture model was established that phenocopied GAMs that predominate in tumors that contact the V-SVZ neural stem cell niche. These GAMs were originally described to have M2-like suppressive features, a finding that was confirmed here. Assessment of GBM secreted proteins identified IL-8 as a key factor necessary for instructing this macrophage identity in GBM.

Upon initial assessment of marker expression of polarized macrophages, we anticipated that a factor such as IL-6 or IL-4 would be secreted across GBM samples, as these cytokines have been well characterized for their role in instructing macrophages towards M2-like suppressive identities^62,63^. In addition, when assessing the protein expression of markers in the M_IL-6 and M_GBM_TCM macrophage phenotypes, we noticed that these two classes shared the expression of some key surface markers (Figure 1A). However, we unexpectedly identified IL-8 as the most robustly secreted molecule among the 105 different soluble proteins that were tested (Figure 2), and we observed that IL-6 was no better than IFNγ and far worse that IL-8 and TCM at producing macrophages like those observed *in vivo*.

Cytokines like IL-6 and IL-8 have been identified as a part of the molecular signature of cytokines secreted by GBM tumors ^64,65^. Yet so far, literature surrounding IL-8 in cancer has focused on its role in neutrophil recruitment^66^ and angiogenesis^67^ during tissue remodeling and inflammation^68^. In a healthy wound healing response setting, activated macrophages, endothelial cells, and epithelial cells can produce IL-8 in response to infections or tissue injury. IL-8 can function as a chemoattractant for neutrophils who may form extracellular traps to kill invading microbes^69^, and in endothelial cells IL-8 signaling can induce cell proliferation, survival and migration leading to angiogenesis ^70^. IL-8 expression has been previously detected and linked to tumor progression in several types of cancers including breast, colon, ovarian, bladder, and prostate cancers, as well as in melanoma^71–74^. In GBM especially, studies have focused on describing the role of IL-8 in promoting glioblastoma stem cells^48^, which are also thought to be major drivers of aggressive GBM tumors, cell migration signaling ^75^, and angiogenesis^46,50,76^. Preclinical studies have suggested that blocking antibody treatments against IL-8 could potentiate greater efficacy of immunotherapies in GBM^77^. While GBM secreted IL-8 has been shown to be an important player driving GBM cancer cell behavior^52^, a major gap in the field remained when it came to understanding how its presence in the TME could be affecting infiltrating macrophages. In fact, when looking further to understand how IL-8 affects macrophages in a healthy tissue remodeling setting, we found that thus far IL-8 has solely been described as a cytokine that is secreted by macrophages, and the features of a macrophage response to this cytokine remained largely unknown until this study. Here we show that IL-8 is secreted both *in vivo* and *ex vivo* by GBM tumor cells, and we show that it is necessary for polarization of M_GBM_TCM macrophages. Ultimately, these results indicate that IL-8 should be further studied as a key determinant of the microenvironment in GBM and other cancer types, as IL-8 and its receptors may represent suitable targets for therapies.

While this study describes the surface protein expression of key markers activated by IL-8 and TCM in polarized macrophages, further studies are necessary to continue to fully describe the impact of IL-8 on polarization of healthy microglia, monocytes and macrophages. Future studies of M_IL-8 polarization should assess functional tests such as antigen presentation, T cell activation/suppression in a mixed leukocyte reaction, and cytokine secretion. In addition, phospho-flow cytometry could be used to dissect downstream signaling pathways that are mechanistically involved in M_IL-8 polarization.

It is well established that immortalized cancer cell lines, which are often used as surrogates for human tumors, provide a poor reflection of the diverse profiles of human patients’ tumors^78^. GBM tumors are especially notorious for their extensive genetic, epigenetic, intratumoral and intertumoral heterogeneity^79^ which is often depicted as one of the major challenges for understanding the cellular and molecular underpinnings driving GBM. It has been shown that the use of patient derived primary tumor cells can provide a model that more closely reproduces the *in vivo* human tumor microenvironment^80,81^. In attempt to understand GBM heterogeneity, transcriptomic analysis of primary tumor samples was recently used to describe 4 distinct transcriptional states: neural progenitor (NPC)-, oligodendrocyte progenitor (OPC)-, astrocytic (AC)-, and mesenchymal (MES)-like ^82^. Other models such as the use of organoids and patient derived xenograft models have also been very useful tools for dissecting GBM tumor heterogeneity and are increasingly utilized in neuro-oncology research for preclinical studies ^83,84^. In this study, the use of primary human GBM tumor cells and healthy blood monocytes provided a model that yielded results with in vivo human relevance. Given that even immune competent mouse models don’t express an IL-8 equivalent, the ex-vivo culture of primary cells provided an appropriate and representative model in this study.

This work suggests a new mechanism for IL-8 in glioblastoma and implicates CD32^+^ macrophages as a key feature of the aggressive, immunosuppressive immune microenvironment in GBM. Clinical studies should now explore targeting of IL-8, especially for patients with V-SVZ contacting GBM.

## Methods

### Tissue Collection of Human Specimens and Processing

Surgical resection specimens of IDH-wild type glioblastomas that were collected at Vanderbilt University Medical Center between 2014 and 2023 were processed into single cell suspensions following an established dissociation protocol^85^. All samples were collected with patient informed consent in compliance with the Vanderbilt Institutional Review Board (IRB #131870), and in accordance with the declaration of Helsinki.

Patients were adults (≥18 years of age) at the time of surgical resection. Resections were classified as gross or subtotal resections by a neurosurgeon and a neuroradiologist. Tumor contact status in relation to the lateral ventricle (V-SVZ) was determined by a neurosurgeon and a radiologist based on pre-operative radiographic magnetic resonance imaging (MRI) of the brain, as detailed in Mistry et al. ^86,87^.

### Ex vivo culture of GBM tumor cells and generation of tumor conditioned media

Cryopreserved samples of single cell dissociated GBM tumor cells were cultured for 3 days in ultra-low attachment plates at a density of 2×10^5^ cells/ml in a humidified atmosphere at 37°C, 5% CO2 in macrophage polarization medium (RPMI 1640 enriched with FBS 10% and supplemented with 1% PenStrep solution^37,88^). The resulting tumor conditioned medium was removed from the cells via centrifugation and stored in 500 µl aliquots at -20 °C for future experiments.

### Macrophage polarization

Peripheral blood mononuclear cells (PBMCs) from healthy donors were obtained commercially (Stem Cell Technologies). For in vitro macrophage polarization experiments, healthy macrophages were obtained by differentiating healthy monocytes. For *ex vivo* differentiation of monocytes, cells were cultured in 6-well plates at 2 × 10^6^ cells/ml in a humidified atmosphere at 37°C, 5% CO2 in RPMI 1640 enriched with FBS 10% and supplemented with 1% Pen/Strep solution. To purify monocytes from PBMCs via plate adhesion, PBMCs were allowed to adhere to the plate for 3 hours to enrich monocytes; the cells in suspension were discarded. To activate macrophage differentiation, monocytes were stimulated with macrophage colony stimulating factor (M-CSF) (50 ng/ml) for 3 d in standard tissue culture, as previously described^34,37^. Then, macrophages were further polarized for 3 d by IL-6 (10 ng/ml), IFN-g (10 ng/ml), or a mixture composed of 50% tumor conditioned media (GBM_TCM) (see TCM section) and 50% fresh media supplemented with M-CSF. At the end of the polarization, wells were treated with Accutase (Sigma-Aldrich) prewarmed at 37°C for 30 s before collection of cells for further analysis.

### IL-8 blockade in ex vivo studies

To block IL-8 during macrophage polarization, a monoclonal antibody against IL-8(ɑ-IL-8) (clone #: 6217) was used. Aliquots of tumor conditioned media were thawed and incubated with 0.8µg/mL of ɑ-IL-8 for 30 minutes at 37°C prior macrophage stimulation. TCM was then combined with complete macrophage media and added to differentiated macrophages to allow for polarization in culture over the next 3 days. Macrophages were then collected accordingly with other conditions for analysis via spectral flow cytometry.

### Spectral flow cytometry

Polarized macrophages were collected post Accutase treatment and transferred to FACS tubes for live staining. Cells were washed once with PBS and once with PBS BSA. Cells were stained for 30 minutes with an antibody cocktail which included CD86 (IT2.2), CD64 (10.1),CD206 (EPR6828(B)), HLA-DR (I243), PD-L1 (29E.2A3), CD163 (GHI/61), CD32 (FUN-2). During the last 5 minutes of staining, the viability dye Alexa 700 SE dye (NHS) was added. Cells were washed once with PBS and once with PBS BSA. After staining was complete, cells were analyzed on a CyTEK spectral flow cytometer within 30-60 minutes.

### Cytokine arrays

The Proteome Profiler Human XL Cytokine Array Kit (R&D Systems) was used to measure the secretion of 105 different soluble factors and cytokines in GBM tumor conditioned media. 500µl of tumor conditioned media from each patient sample were analyzed per the manufacturer’s instructions on a nitrocellulose membrane and then visualized using chemiluminescent detection reagents and the iBright imaging system.

### Human tumor microarray

Generation of a tumor microarray (TMA) of formalin fixed paraffin embedded (FFPE) glioblastoma specimens was described in Leelatian et. al. ^89^. Briefly, three 1mm areas were selected from each tumor sample by a neuropathologist. Blocks were delivered to the Vanderbilt University Medical Center TPSR (Translational Pathology Shared Resource), where cores were extracted from the encircled areas using the Tissue Microarray Grandmaster (3DHistech). IHC of serial sections of the two resulting TMA blocks (<10 μm thick) were stained with primary antibodies conjugated to HRP and 3,3′-Diaminobenzidine (DAB) detection for CXCL8 (clone), and counter stained with hematoxylin by the TPSR. Digital images were obtained with an Ariol SL-50 automated scanning microscope and the Leica SCN400 Slide Scanner from VUMC Digital Histology Shared Resource.

### Data processing

All flow cytometry data collected was uploaded to Cytobank for further analysis.

TMA IHC data was de-arrayed using QuPath (version 0.5.0) and per pixel DAB signal was quantified in python using the separate_stains function in scikit-image (version 0.22.0). Individual core images were processed for display using ImageJ.

For cytokine arrays, ImageJ Dot blot analyzer macro^90^ was used to quantify intensity density of dots present on each dot blot.

## Data Availability

Datasets analyzed in this manuscript are online at FlowRepository^91^ made available for reviewers (see link in submission materials), and will be made public upon acceptance. Transparent analysis scripts for datasets in this manuscript are available on the CytoLab Github page (https://github.com/cytolab/) with open-source code and commented Rmarkdown analysis walkthroughs.

## Acknowledgements

We thank Vanderbilt’s Cancer and Immunology Core and Flow Cytometry Shared Resource facilities as well as all the surgeons, patients, and families that supported this work. Research was supported by the following funding resources: NIH/NCI grants R01 NS096238 (RAI, JMI), R01 CA226833 (JMI, SM, CEC, MJH), R01 NS118580 (RAI, AAB), U01 AI125056 (JMI), U54 CA217450 (JMI, MJH), T32 GM139800 (SM), T32 AI138932 (SM), the Vanderbilt-Ingram Cancer Center (VICC, P30 CA68485), the Michael David Greene Brain Cancer Fund (RAI, JMI), the Southeastern Brain Tumor Foundation (RAI, JMI), a gift from Daniel F Hewins (RAI), and the Ben & Catherine Ivy Foundation (RAI, JMI). Translational Pathology Shared Resource (TPSR) is supported by NCI/NIH Cancer Center Support Grant P30CA068485. We thank outstanding undergraduate students Alejandra Rosario-Crespo, Amanda Kouaho, and Niraj Rama for their work on projects exploring cancer and immune cell identity.

## Author Contributions

SM, RAI, and JMI designed and conceptualized the study. RAI, and JMI, provided intellectual support in assembling datasets. SM, MJH, and AAB collected data. SM, CEC, and AAB developed data analysis scripts. SM and JMI performed flow cytometry data analysis and interpretation. AAM and SC scored MRI images for tumor contact with the lateral ventricle and provided patients’ clinical characteristics. BCM confirmed tissue pathology. LBC, RCT, and KDW provided freshly resected tissue specimens. SM and JMI wrote the manuscript. RAI and JMI provided financial support. All authors contributed in reviewing and editing the manuscript.

## References

1. Davis, M. (2016). Glioblastoma: Overview of Disease and Treatment. Clinical Journal of Oncology Nursing 20, S2–S8. 10.1188/16.cjon.s1.2-8.

2. Ostrom, Q.T., Cioffi, G., Gittleman, H., Patil, N., Waite, K., Kruchko, C., and Barnholtz-Sloan, J.S. (2019). CBTRUS Statistical Report: Primary Brain and Other Central Nervous System Tumors Diagnosed in the United States in 2012–2016. Neuro-Oncology 21, v1–v100. 10.1093/neuonc/noz150.

3. Sheikh, S., Radivoyevitch, T., Barnholtz-Sloan, J.S., and Vogelbaum, M. (2019). Long-term trends in glioblastoma survival: implications for historical control groups in clinical trials. Neuro-Oncology Practice. 10.1093/nop/npz046.

4. Zhang, Y., and Zhang, Z. (2020). The history and advances in cancer immunotherapy: understanding the characteristics of tumor-infiltrating immune cells and their therapeutic implications. Cellular & Molecular Immunology 17, 807–821. 10.1038/s41423-020-0488-6.

5. Fecci, P.E., and Sampson, J.H. (2019). The current state of immunotherapy for gliomas: an eye toward the future. Journal of Neurosurgery 131, 657–666. 10.3171/2019.5.jns181762.

6. Fu, W., Wang, W., Li, H., Jiao, Y., Huo, R., Yan, Z., Wang, J., Wang, S., Chen, D., Cao, Y., and Zhao, J. Single-Cell Atlas Reveals Complexity of the Immunosuppressive Microenvironment of Initial and Recurrent Glioblastoma.

7. Gutmann, D.H., and Kettenmann, H. (2019). Microglia/Brain Macrophages as Central Drivers of Brain Tumor Pathobiology. Neuron 104, 442–449. 10.1016/j.neuron.2019.08.028.

8. Yao, Y., Ye, H., Qi, Z., Mo, L., Yue, Q., Baral, A., Hoon, D.S.B., Vera, J.C., Heiss, J.D., Chen, C.C., et al. (2016). B7-H4(B7x)-Mediated Cross-talk between Glioma-Initiating Cells and Macrophages via the IL6/JAK/STAT3 Pathway Lead to Poor Prognosis in Glioma Patients. Clin Cancer Res 22, 2778–2790. 10.1158/1078-0432.ccr-15-0858.

9. Khan, F., Pang, L., Dunterman, M., Lesniak, M.S., Heimberger, A.B., and Chen, P. (2023). Macrophages and microglia in glioblastoma: heterogeneity, plasticity, and therapy. Journal of Clinical Investigation 133. 10.1172/jci163446.

10. Hambardzumyan, D., Gutmann, D.H., and Kettenmann, H. (2016). The role of microglia and macrophages in glioma maintenance and progression. Nat Neurosci 19, 20–27. 10.1038/nn.4185.

11. Chen, Z., Feng, X., Herting, C.J., Garcia, V.A., Nie, K., Pong, W.W., Rasmussen, R., Dwivedi, B., Seby, S., Wolf, S.A., et al. Cellular and Molecular Identity of Tumor-Associated Macrophages in Glioblastoma.

12. Mignogna, C., Signorelli, F., Vismara, M.F., Zeppa, P., Camastra, C., Barni, T., Donato, G., and Di Vito, A. (2016). A reappraisal of macrophage polarization in glioblastoma: Histopathological and immunohistochemical findings and review of the literature. Pathol Res Pract 212, 491–499. 10.1016/j.prp.2016.02.020.

13. Walentynowicz, K.A., Engelhardt, D., Cristea, S., Yadav, S., Onubogu, U., Salatino, R., Maerken, M., Vincentelli, C., Jhaveri, A., Geisberg, J., et al. Single-cell heterogeneity of EGFR and CDK4 co-amplification is linked to immune infiltration in glioblastoma.

14. Liu, S., Zhang, C., Maimela, N.R., Yang, L., Zhang, Z., Ping, Y., Huang, L., and Zhang, Y. Molecular and clinical characterization of CD163 expression via large-scale analysis in glioma.

15. Bartkowiak, T., Lima, S.M., Hayes, M.J., Mistry, A.M., Brockman, A.A., Sinnaeve, J., Leelatian, N., Roe, C.E., Mobley, B.C., Chotai, S., et al. (2023). An immunosuppressed microenvironment distinguishes lateral ventricle–contacting glioblastomas. JCI Insight 8. 10.1172/jci.insight.160652.

16. Mistry, A.M., Hale, A.T., Chambless, L.B., Weaver, K.D., Thompson, R.C., and Ihrie, R.A. (2017). Influence of glioblastoma contact with the lateral ventricle on survival: a meta-analysis. J Neurooncol 131, 125–133. 10.1007/s11060-016-2278-7.

17. Mistry, A.M., Dewan, M.C., White-Dzuro, G.A., Brinson, P.R., Weaver, K.D., Thompson, R.C., Ihrie, R.A., and Chambless, L.B. Decreased survival in glioblastomas is specific to contact with the ventricular-subventricular zone, not subgranular zone or corpus callosum.

18. Poon, C.C., Sarkar, S., Yong, V.W., and Kelly, J.J.P. (2017). Glioblastoma-associated microglia and macrophages: targets for therapies to improve prognosis. Brain 140, 1548–1560. 10.1093/brain/aww355.

19. Ye, Z., Ai, X., Yang, K., Yang, Z., Fei, F., Liao, X., Qiu, Z., Gimple, R.C., Yuan, H., Huang, H., et al. (2023). Targeting Microglial Metabolic Rewiring Synergizes with Immune-Checkpoint Blockade Therapy for Glioblastoma. Cancer Discov 13, 974–1001. 10.1158/2159-8290.Cd-22-0455.

20. Fermi, V., Warta, R., Wöllner, A., Lotsch, C., Jassowicz, L., Rapp, C., Knoll, M., Jungwirth, G., Jungk, C., Dao Trong, P., et al. (2023). Effective Reprogramming of Patient-Derived M2-Polarized Glioblastoma-Associated Microglia/Macrophages by Treatment with GW2580. Clin Cancer Res 29, 4685–4697. 10.1158/1078-0432.Ccr-23-0576.

21. Quail, D.F., and Joyce, J.A. Microenvironmental regulation of tumor progression and metastasis.

22. Zhang, Y., Cheng, S., Zhang, M., Zhen, L., Pang, D., Zhang, Q., and Li, Z. (2013). High-infiltration of tumor-associated macrophages predicts unfavorable clinical outcome for node-negative breast cancer. PLoS One 8, e76147. 10.1371/journal.pone.0076147.

23. Kumar, A.T., Knops, A., Swendseid, B., Martinez-Outschoom, U., Harshyne, L., Philp, N., Rodeck, U., Luginbuhl, A., Cognetti, D., Johnson, J., and Curry, J. (2019). Prognostic Significance of Tumor-Associated Macrophage Content in Head and Neck Squamous Cell Carcinoma: A Meta-Analysis. Front Oncol 9, 656. 10.3389/fonc.2019.00656.

24. Koll, F.J., Banek, S., Kluth, L., Köllermann, J., Bankov, K., Chun, F.K., Wild, P.J., Weigert, A., and Reis, H. (2023). Tumor-associated macrophages and Tregs influence and represent immune cell infiltration of muscle-invasive bladder cancer and predict prognosis. J Transl Med 21, 124. 10.1186/s12967-023-03949-3.

25. Wang, H., Yang, L., Wang, D., Zhang, Q., and Zhang, L. (2017). Pro-tumor activities of macrophages in the progression of melanoma. Hum Vaccin Immunother 13, 1556–1562. 10.1080/21645515.2017.1312043.

26. Erlandsson, A., Carlsson, J., Lundholm, M., Fält, A., Andersson, S.-O., Andrén, O., and Davidsson, S. (2019). M2 macrophages and regulatory T cells in lethal prostate cancer. The Prostate 79, 363–369. 10.1002/pros.23742.

27. Szulzewsky, F., Pelz, A., Feng, X., Synowitz, M., Markovic, D., Langmann, T., Holtman, I.R., Wang, X., Eggen, B.J., Boddeke, H.W., et al. (2015). Glioma-associated microglia/macrophages display an expression profile different from M1 and M2 polarization and highly express Gpnmb and Spp1. PLoS One 10, e0116644. 10.1371/journal.pone.0116644.

28. Chen, D., Varanasi, S.K., Hara, T., Traina, K., Sun, M., McDonald, B., Farsakoglu, Y., Clanton, J., Xu, S., Garcia-Rivera, L., et al. CTLA-4 blockade induces a microglia-Th1 cell partnership that stimulates microglia phagocytosis and anti-tumor function in glioblastoma.

29. Gangoso, E., Southgate, B., Bradley, L., Rus, S., Galvez-Cancino, F., McGivern, N., Güç, E., Kapourani, C.-A., Byron, A., Ferguson, K.M., et al. (2021). Glioblastomas acquire myeloid-affiliated transcriptional programs via epigenetic immunoediting to elicit immune evasion. Cell 184, 2454–2470.e2426. 10.1016/j.cell.2021.03.023.

30. Yang, F.A.-O., Akhtar, M.A.-O., Zhang, D.A.-O., El-Mayta, R.A.-O.X., Shin, J.A.-O., Dorsey, J.F., Zhang, L., Xu, X.A.-O., Guo, W., Bagley, S.A.-O., et al. An immunosuppressive vascular niche drives macrophage polarization and immunotherapy resistance in glioblastoma.

31. Chryplewicz, A., Scotton, J., Tichet, M., Zomer, A., Shchors, K., Joyce, J.A., Homicsko, K., and Hanahan, D. Cancer cell autophagy, reprogrammed macrophages, and remodeled vasculature in glioblastoma triggers tumor immunity.

32. Murray, J., Peter, Allen, E., Judith, Biswas, K., Subhra, Fisher, A., Edward, Gilroy, W., Derek, Goerdt, S., Gordon, S., Hamilton, A., John, Ivashkiv, B., Lionel, Lawrence, T., et al. (2014). Macrophage Activation and Polarization: Nomenclature and Experimental Guidelines. Immunity 41, 14–20. 10.1016/j.immuni.2014.06.008.

33. Roussel, M., Ferrell, P.B., Greenplate, A.R., Lhomme, F., Le Gallou, S., Diggins, K.E., Johnson, D.B., and Irish, J.M. (2017). Mass cytometry deep phenotyping of human mononuclear phagocytes and myeloid-derived suppressor cells from human blood and bone marrow. Journal of Leukocyte Biology 102, 437–447. 10.1189/jlb.5ma1116-457r.

34. Xue, J., Schmidt, V., Susanne, Sander, J., Draffehn, A., Krebs, W., Quester, I., Nardo, D., Dominic, Gohel, D., Trupti, Emde, M., Schmidleithner, L., et al. (2014). Transcriptome-Based Network Analysis Reveals a Spectrum Model of Human Macrophage Activation. Immunity 40, 274–288. 10.1016/j.immuni.2014.01.006.

35. Chen, R.A.-O., Yang, D., Shen, L., Fang, J., Khan, R., and Liu, D.A.-O. Overexpression of CD86 enhances the ability of THP-1 macrophages to defend against Talaromyces marneffei.

36. Schulz, D., Severin, Y., Zanotelli, V.R.T., and Bodenmiller, B. In-Depth Characterization of Monocyte-Derived Macrophages using a Mass Cytometry-Based Phagocytosis Assay.

37. Roussel, M., Bartkowiak, T., and Irish, J.M. (2019). Picturing Polarized Myeloid Phagocytes and Regulatory Cells by Mass Cytometry. Methods in molecular biology 1989, 217–226. 10.1007/978-1-4939-9454-0_14.

38. Peter, Judith, Subhra, Edward, Derek, Goerdt, S., Gordon, S., John, Lionel, Lawrence, T., et al. (2014). Macrophage Activation and Polarization: Nomenclature and Experimental Guidelines. Immunity 41, 14–20. 10.1016/j.immuni.2014.06.008.

39. Bronte, V., Brandau, S., Chen, S.-H., Colombo, M.P., Frey, A.B., Greten, T.F., Mandruzzato, S., Murray, P.J., Ochoa, A., Ostrand-Rosenberg, S., et al. (2016). Recommendations for myeloid-derived suppressor cell nomenclature and characterization standards. Nature Communications 7, 12150. 10.1038/ncomms12150.

40. Sharma, I., Singh, A., Sharma, K., and Saxena, S. (2017). Gene Expression Profiling of Chemokines and Their Receptors in Low and High Grade Astrocytoma. Asian Pac J Cancer Prev 18, 1307–1313. 10.22034/apjcp.2017.18.5.1307.

41. Raghuwanshi, S.K., Su, Y., Singh, V., Haynes, K., Richmond, A., and Richardson, R.M. (2012). The Chemokine Receptors CXCR1 and CXCR2 Couple to Distinct G Protein-Coupled Receptor Kinases To Mediate and Regulate Leukocyte Functions. The Journal of Immunology 189, 2824–2832. 10.4049/jimmunol.1201114.

42. Holmes, W.E., Lee, J., Kuang, W.J., Rice, G.C., and Wood, W.I. (1991). Structure and functional expression of a human interleukin-8 receptor. Science 253, 1278–1280. 10.1126/science.1840701.

43. Waugh, D.J.J., and Wilson, C. (2008). The Interleukin-8 Pathway in Cancer. Clinical Cancer Research 14, 6735–6741. 10.1158/1078-0432.Ccr-07-4843.

44. Heidemann, J., Ogawa, H., Dwinell, M.B., Rafiee, P., Maaser, C., Gockel, H.R., Otterson, M.F., Ota, D.M., Lugering, N., Domschke, W., and Binion, D.G. (2003). Angiogenic effects of interleukin 8 (CXCL8) in human intestinal microvascular endothelial cells are mediated by CXCR2. J Biol Chem 278, 8508–8515. 10.1074/jbc.M208231200.

45. Prince, E.W., Apps, J.R., Jeang, J., Chee, K., Medlin, S., Jackson, E.M., Dudley, R., Limbrick, D., Naftel, R., Johnston, J., et al. (2024). Unraveling the Complexity of the Senescence-Associated Secretory Phenotype in Adamantinomatous Craniopharyngioma Using Multi-Modal Machine Learning Analysis. Neuro-Oncology, noae015. 10.1093/neuonc/noae015.

46. Chen, Z., Will, R., Kim, S.N., Busch, M.A., Dünker, N., Dammann, P., Sure, U., and Zhu, Y. (2023). Novel Function of Cancer Stem Cell Marker ALDH1A3 in Glioblastoma: Pro-Angiogenesis through Paracrine PAI-1 and IL-8. Cancers 15, 4422. 10.3390/cancers15174422.

47. Brat, D.J., Bellail, A.C., and Van Meir, E.G. (2005). The role of interleukin-8 and its receptors in gliomagenesis and tumoral angiogenesis. Neuro Oncol 7, 122–133. 10.1215/s1152851704001061.

48. Infanger, D.W., Cho, Y., Lopez, B.S., Mohanan, S., Liu, S.C., Gursel, D., Boockvar, J.A., and Fischbach, C. (2013). Glioblastoma stem cells are regulated by interleukin-8 signaling in a tumoral perivascular niche. Cancer Res 73, 7079–7089. 10.1158/0008-5472.Can-13-1355.

49. Sharma, I., Singh, A., Siraj, F., and Saxena, S. (2018). IL-8/CXCR1/2 signalling promotes tumor cell proliferation, invasion and vascular mimicry in glioblastoma. J Biomed Sci 25, 62. 10.1186/s12929-018-0464-y.

50. Hasan, T., Caragher, S.P., Shireman, J.M., Park, C.H., Atashi, F., Baisiwala, S., Lee, G., Guo, D., Wang, J.Y., Dey, M., et al. (2019). Interleukin-8/CXCR2 signaling regulates therapy-induced plasticity and enhances tumorigenicity in glioblastoma. Cell Death & Disease 10, 292. 10.1038/s41419-019-1387-6.

51. Hasan, T., Caragher, S.P., Shireman, J.M., Park, C.H., Atashi, F., Baisiwala, S., Lee, G., Guo, D., Wang, J.Y., Dey, M., et al. (2019). Interleukin-8/CXCR2 signaling regulates therapy-induced plasticity and enhances tumorigenicity in glioblastoma. Cell Death & Disease 10. 10.1038/s41419-019-1387-6.

52. Holst, C.B., Christensen, I.J., Vitting-Seerup, K., Skjøth-Rasmussen, J., Hamerlik, P., Poulsen, H.S., and Johansen, J.S. (2021). Plasma IL-8 and ICOSLG as prognostic biomarkers in glioblastoma. Neurooncol Adv 3, vdab072. 10.1093/noajnl/vdab072.

53. Benner, B., Scarberry, L., Suarez-Kelly, L.P., Duggan, M.C., Campbell, A.R., Smith, E., Lapurga, G., Jiang, K., Butchar, J.P., Tridandapani, S., et al. (2019). Generation of monocyte-derived tumor-associated macrophages using tumor-conditioned media provides a novel method to study tumor-associated macrophages in vitro. Journal for ImmunoTherapy of Cancer 7. 10.1186/s40425-019-0622-0.

54. Diggins, K.E., Greenplate, A.R., Leelatian, N., Wogsland, C.E., and Irish, J.M. (2017). Characterizing cell subsets using marker enrichment modeling. Nature Methods 14, 275–278. 10.1038/nmeth.4149.

55. Greenplate, A.R., McClanahan, D.D., Oberholtzer, B.K., Doxie, D.B., Roe, C.E., Diggins, K.E., Leelatian, N., Rasmussen, M.L., Kelley, M.C., Gama, V., et al. (2019). Computational Immune Monitoring Reveals Abnormal Double-Negative T Cells Present across Human Tumor Types. Cancer Immunol Res 7, 86–99. 10.1158/2326-6066.CIR-17-0692.

56. Barone, S.M., Paul, A.G., Muehling, L.M., Lannigan, J.A., Kwok, W.W., Turner, R.B., Woodfolk, J.A., and Irish, J.M. (2021). Unsupervised machine learning reveals key immune cell subsets in COVID-19, rhinovirus infection, and cancer therapy. eLife 10. 10.7554/elife.64653.

57. Hanahan, D., and Robert (2011). Hallmarks of Cancer: The Next Generation. Cell 144, 646–674. 10.1016/j.cell.2011.02.013.

58. Chen, Daniel S., and Mellman, I. (2013). Oncology Meets Immunology: The Cancer-Immunity Cycle. Immunity 39, 1–10. 10.1016/j.immuni.2013.07.012.

59. Nørøxe, D.S., Poulsen, H.S., and Lassen, U. (2016). Hallmarks of glioblastoma: a systematic review. ESMO open. 1, e000144. 10.1136/esmoopen-2016-000144.

60. Martinez-Lage, M., Lynch, T.M., Bi, Y., Cocito, C., Way, G.P., Pal, S., Haller, J., Yan, R.E., Ziober, A., Nguyen, A., et al. (2019). Immune landscapes associated with different glioblastoma molecular subtypes. Acta Neuropathologica Communications 7. 10.1186/s40478-019-0803-6.

61. Razavi, S.M., Lee, K.E., Jin, B.E., Aujla, P.S., Gholamin, S., and Li, G. (2016). Immune Evasion Strategies of Glioblastoma. Front Surg 3, 11. 10.3389/fsurg.2016.00011.

62. Wang, H.W., and Joyce, J.A. (2010). Alternative activation of tumor-associated macrophages by IL-4: priming for protumoral functions. Cell Cycle 9, 4824–4835. 10.4161/cc.9.24.14322.

63. Shapouri-Moghaddam, A., Mohammadian, S., Vazini, H., Taghadosi, M., Esmaeili, S.A., Mardani, F., Seifi, B., Mohammadi, A., Afshari, J.T., and Sahebkar, A. (2018). Macrophage plasticity, polarization, and function in health and disease. J Cell Physiol 233, 6425–6440. 10.1002/jcp.26429.

64. Tobin, R.P., Jordan, K.R., Kapoor, P., Spongberg, E., Davis, D., Vorwald, V.M., Couts, K.L., Gao, D., Smith, D.E., Borgers, J.S.W., et al. (2019). IL-6 and IL-8 Are Linked With Myeloid-Derived Suppressor Cell Accumulation and Correlate With Poor Clinical Outcomes in Melanoma Patients. Frontiers in Oncology 9.

65. Jarmuzek, P., Defort, P., Kot, M., Wawrzyniak-Gramacka, E., Morawin, B., and Zembron-Lacny, A. (2023). Cytokine Profile in Development of Glioblastoma in Relation to Healthy Individuals. Int J Mol Sci 24. 10.3390/ijms242216206.

66. Baggiolini, M., Walz, A., and Kunkel, S.L. (1989). Neutrophil-activating peptide-1/interleukin 8, a novel cytokine that activates neutrophils. J Clin Invest 84, 1045–1049. 10.1172/jci114265.

67. Li, A., Dubey, S., Varney, M.L., Dave, B.J., and Singh, R.K. (2003). IL-8 directly enhanced endothelial cell survival, proliferation, and matrix metalloproteinases production and regulated angiogenesis. J Immunol 170, 3369–3376. 10.4049/jimmunol.170.6.3369.

68. David, J.M., Dominguez, C., Hamilton, D.H., and Palena, C. (2016). The IL-8/IL-8R Axis: A Double Agent in Tumor Immune Resistance. Vaccines (Basel) 4. 10.3390/vaccines4030022.

69. Brinkmann, V., Reichard, U., Goosmann, C., Fauler, B., Uhlemann, Y., Weiss, D.S., Weinrauch, Y., and Zychlinsky, A. (2004). Neutrophil extracellular traps kill bacteria. Science 303, 1532–1535. 10.1126/science.1092385.

70. Lattanzio, L., Tonissi, F., Torta, I., Gianello, L., Russi, E., Milano, G., Merlano, M., and Lo Nigro, C. (2013). Role of IL-8 induced angiogenesis in uveal melanoma. Invest New Drugs 31, 1107–1114. 10.1007/s10637-013-0005-1.

71. Xie, K. (2001). Interleukin-8 and human cancer biology. Cytokine & growth factor reviews. 12, 375–391. 10.1016/S1359-6101(01)00016-8.

72. Filimon, A., Preda, I.A., Boloca, A.F., and Negroiu, G. (2021). Interleukin-8 in Melanoma Pathogenesis, Prognosis and Therapy-An Integrated View into Other Neoplasms and Chemokine Networks. Cells 11. 10.3390/cells11010120.

73. Zhang, W., Yang, F., Zheng, Z., Li, C., Mao, S., Wu, Y., Wang, R., Zhang, J., Zhang, Y., Wang, H., et al. (2022). Sulfatase 2 Affects Polarization of M2 Macrophages through the IL-8/JAK2/STAT3 Pathway in Bladder Cancer. Cancers 15, 131. 10.3390/cancers15010131.

74. Aalinkeel, R., Nair, B., Chen, C.K., Mahajan, S.D., Reynolds, J.L., Zhang, H., Sun, H., Sykes, D.E., Chadha, K.C., Turowski, S.G., et al. (2016). Nanotherapy silencing the interleukin-8 gene produces regression of prostate cancer by inhibition of angiogenesis. Immunology 148, 387–406. 10.1111/imm.12618.

75. Zhang, B., Shi, L., Lu, S., Sun, X., Liu, Y., Li, H., Wang, X., Zhao, C., Zhang, H., and Wang, Y. (2015). Autocrine IL-8 promotes F-actin polymerization and mediate mesenchymal transition via ELMO1-NF-κB-Snail signaling in glioma. Cancer Biol Ther 16, 898–911. 10.1080/15384047.2015.1028702.

76. Dumitru, C.A., Schröder, H., Schäfer, F.T.A., Aust, J.F., Kreße, N., Siebert, C.L.R., Stein, K.-P., Haghikia, A., Wilkens, L., Mawrin, C., and Sandalcioglu, I.E. (2023). Progesterone Receptor Membrane Component 1 (PGRMC1) Modulates Tumour Progression, the Immune Microenvironment and the Response to Therapy in Glioblastoma. Cells 12, 2498. 10.3390/cells12202498.

77. Liu, H., Zhao, Q., Tan, L., Wu, X., Huang, R., Zuo, Y., Chen, L., Yang, J., Zhang, Z.-X., Ruan, W., et al. (2023). Neutralizing IL-8 potentiates immune checkpoint blockade efficacy for glioma. Cancer cell. 41, 693–710.e698. 10.1016/j.ccell.2023.03.004.

78. Ross, D.T., and Perou, C.M. (2001). A comparison of gene expression signatures from breast tumors and breast tissue derived cell lines. Dis Markers 17, 99–109. 10.1155/2001/850531.

79. Vartanian, A., Singh, S.K., Agnihotri, S., Jalali, S., Burrell, K., Aldape, K.D., and Zadeh, G. (2014). GBM’s multifaceted landscape: highlighting regional and microenvironmental heterogeneity. Neuro-Oncology 16, 1167–1175. 10.1093/neuonc/nou035.

80. Leblanc, V.G., Trinh, D.L., Aslanpour, S., Hughes, M., Livingstone, D., Jin, D., Ahn, B.Y., Blough, M.D., Cairncross, J.G., Chan, J.A., et al. (2022). Single-cell landscapes of primary glioblastomas and matched explants and cell lines show variable retention of inter- and intratumor heterogeneity. Cancer Cell 40, 379–392.e379. 10.1016/j.ccell.2022.02.016.

81. Lum, D.H., Matsen, C., Welm, A.L., and Welm, B.E. (2012). Overview of human primary tumorgraft models: comparisons with traditional oncology preclinical models and the clinical relevance and utility of primary tumorgrafts in basic and translational oncology research. Curr Protoc Pharmacol Chapter 14, Unit 14.22. 10.1002/0471141755.ph1422s59.

82. Neftel, C., Laffy, J., Filbin, M.G., Hara, T., Shore, M.E., Rahme, G.J., Richman, A.R., Silverbush, D., Shaw, M.L., Hebert, C.M., et al. (2019). An Integrative Model of Cellular States, Plasticity, and Genetics for Glioblastoma. Cell 178, 835–849.e821. 10.1016/j.cell.2019.06.024.

83. Jacob, F., Salinas, R.D., Zhang, D.Y., Nguyen, P.T.T., Schnoll, J.G., Wong, S.Z.H., Thokala, R., Sheikh, S., Saxena, D., Prokop, S., et al. (2020). A Patient-Derived Glioblastoma Organoid Model and Biobank Recapitulates Inter- and Intra-tumoral Heterogeneity. Cell 180, 188–204.e122. 10.1016/j.cell.2019.11.036.

84. Vaubel, R.A., Tian, S., Remonde, D., Schroeder, M.A., Mladek, A.C., Kitange, G.J., Caron, A., Kollmeyer, T.M., Grove, R., Peng, S., et al. (2020). Genomic and Phenotypic Characterization of a Broad Panel of Patient-Derived Xenografts Reflects the Diversity of Glioblastoma. Clin Cancer Res 26, 1094–1104. 10.1158/1078-0432.Ccr-19-0909.

85. Leelatian, N., Doxie, D.B., Greenplate, A.R., Sinnaeve, J., Ihrie, R.A., and Irish, J.M. Preparing Viable Single Cells from Human Tissue and Tumors for Cytomic Analysis.

86. Mistry, A.M., Dewan, M.C., White-Dzuro, G.A., Brinson, P.R., Weaver, K.D., Thompson, R.C., Ihrie, R.A., and Chambless, L.B. (2017). Decreased survival in glioblastomas is specific to contact with the ventricular-subventricular zone, not subgranular zone or corpus callosum. Journal of Neuro-Oncology 132, 341–349. 10.1007/s11060-017-2374-3.

87. Mistry, A.M., Wooten, D.J., Davis, L.T., Mobley, B.C., Quaranta, V., and Ihrie, R.A. (2019). Ventricular-Subventricular Zone Contact by Glioblastoma is Not Associated with Molecular Signatures in Bulk Tumor Data. Scientific Reports 9, 1842. 10.1038/s41598-018-37734-w.

88. Davies, J.Q., and Gordon, S. Isolation and culture of human macrophages.

89. Leelatian, N., Sinnaeve, J., Mistry, A.M., Barone, S.M., Brockman, A.A., Diggins, K.E., Greenplate, A.R., Weaver, K.D., Thompson, R.C., Chambless, L.B., et al. (2020). Unsupervised machine learning reveals risk stratifying glioblastoma tumor cells. eLife 9. 10.7554/elife.56879.

90. Klemm, A.H. (2020). Semi-automated analysis of dot blots using ImageJ/Fiji. F1000Research 9, 1385. 10.12688/f1000research.27179.1.

91. Spidlen, J., Breuer, K., Rosenberg, C., Kotecha, N., and Brinkman, R.R. (2012). FlowRepository: A resource of annotated flow cytometry datasets associated with peer-reviewed publications. Cytometry Part A 81A, 727–731. 10.1002/cyto.a.22106.

